# Early-Enrichment Hit Discovery via Reversible-Work c(t) Estimation in Metadynamics (CTMD)

**DOI:** 10.64898/2026.02.05.703972

**Authors:** Venkata Sai Sreyas Adury, Pratyush Tiwary, Xinyu Gu, Mrinal Shekhar

## Abstract

Virtual screening for small-molecule binders is often limited by false positives from approximate scoring functions and rigid-receptor assumptions. These can be addressed downstream through accurate but expensive free energy calculations. At the same time, recent artificial-intelligence-based co-folding methods have been proposed that claim to achieve accuracy of free energy methods at much lower cost, but these have not yet delivered consistent improvements in early enrichment and can be confounded by memorization. Here we address this gap by introducing c(t)-based metadynamics (CTMD), a physics-based, high-throughput hit-triaging protocol tailored for early enrichment. CTMD uses the nonequilibrium reversible-work estimator *c*(*t*) introduced by Tiwary and Parrinello (Journal of Physical Chemistry B, 2015 119 736), computed from a small number of short, independent well-tempered metadynamics trajectories, to rank binding stability without requiring converged binding free energies. Across diverse targets and chemotypes, CTMD provides robust early enrichment while remaining fast, transferable with minimal parameter tuning, and resistant to memorization-driven artifacts—underscoring both an immediately deployable physics-based alternative for screening. For these systems we show how co-folding, particularly Boltz-2, achieves enrichment directly proportional to similairty with training set, and more worrying, even in the presence of signficant modifications to active site. Given its simplicity of implementation, CTMD should thus be an “embarassingly” open-source, early enrichment method available for use by the broad pharma and academic community that sits right between approximate but fast docking or AI based co-folding methods, and more expensive but accurate free energy calculations, expected to lead to saving significant financial and human capital in drug discovery campaigns.

## Introduction

Small molecule therapeutics remain a cornerstone of modern drug discovery due to their favorable pharmacokinetic profiles, synthetic accessibility, and cost-effectiveness. ^1,2^ Historically, drug discovery programs have relied on experimental high-throughput screening (HTS)^3^ to identify novel chemical matter. However, HTS campaigns are resource-intensive, requiring costly automation, and frequently suffer from low hit rates. In recent years, virtual screening has emerged as a compelling alternative, bolstered by the availability of make-on-demand libraries such as Enamine REAL.^4^ Nevertheless, despite its potential, virtual screening is hindered by high false-positive rates resulting from inaccurate empirical scoring functions and the oversimplification of receptors as rigid entities.

To address these limitations, two distinct approaches have gained traction. A more recent, very enticing strategy employs AI-based co-folding methods, predominantly AlphaFold-3 and Boltz-2,^5,6^ which aim to directly predict protein-ligand complex structures. Although these methods have shown promise in proteinligand structure prediction, a recent benchmark study by Fraser and co-authors^7^ shows that co-folding methods do not yet demonstrate a statistically significant improvement in enriching binders over non-binders compared to traditional structure-based virtual screening approaches.

An older and widely used strategy involves physics-based methods, which explicitly model molecular interactions using molecular mechanics and statistical thermodynamics. Among these, Absolute Binding Free Energy Perturbation (FEP) calculations represent the gold standard in computational affinity prediction, providing thermodynamically rigorous estimates.^8^ However, the computational cost and the requirement for relatively significant technical expertise limit the routine application of FEP in large-scale virtual screening campaigns. A complementary class of physics-based approaches is provided by enhanced sampling methods such as metadynamics, which explicitly describe ligand binding and unbinding along predefined collective variables. While these methods have been successfully applied to compute binding free energies and kinetics,^9,10^ they suffer from inherent challenges: the need for careful collective variable selection, substantial parameter tuning, convergence uncertainties, and computational costs that scale poorly with library size.

Around a decade ago, Clark et al^11^ came up with a simple approach to implement metadynamics in a high throughput manner for ranking docking poses for a given ligand generated from Induced Fit Docking (IFD). This approach, later called Binding Pose Metady-namics (BPMD) and revisited by many publications^12^ presents a computationally efficient method designed to assess the stability of predicted binding poses for a given ligand. By applying a metadynamics bias that perturbs the ligand away from its initial pose, BPMD quantifies pose stability based on the ligand’s resistance to displacement, without requiring convergence of the full binding free energy surface. The bias is applied as a function of the Root Mean Squared Displacement (RMSD) of the ligand from the starting pose, which is a crude but generic variable that does not need much system-specific hand tuning. While BPMD is useful in drug discovery and has become a standard part of IFD protocols, ^13^ it is fundamentally engineered to evaluate pose stability rather than comparing binding affinities of different candidate hit molecules. Consequently, it is ill-suited as a primary metric for large-scale virtual screening of chemically diverse ligands.

Our motivation is this work is to come up with an effective method for hit triaging in virtual screening. We believe it must satisfy three criteria: it must be 1) accurate yet computationally efficient, 2) generalizable with minimal parameter tuning, and 3) a simple framework easy to implement and use by general practitioners. With these requirements in mind, we introduce CTMD (*c*(*t*)-based metadynamics), a physics-informed framework designed for early-enrichment hit discovery. CTMD uses a reversible work estimator for metadynamics introduced as *c*(*t*) by Tiwary and Parrinello in Ref.^14^ Instead of seeking fully converged binding free energies, the method uses nonequilibrium, reversible work signatures to efficiently discriminate ligand binding stability across diverse chemotypes. The central idea is to calculate the lowest value achieved by the quantity *c*(*t*) over a few short, independent well-tempered metadynamics simulations, which serves as a proxy for the thermodynamic instability for a given ligand-receptor pose. *c*(*t*) is implemented as:^15^

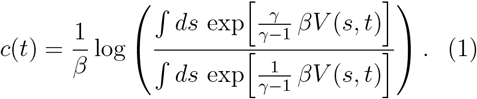

where *s* is the biased collective variable, *β* is the inverse temperature, *t* is the simulation time, *V* (*s, t*) is the time-dependent metadynamics bias, and *γ* is the well-tempered metadynamics bias factor. We keep the same *s* as in BPMD. We provide further technical details later in CTMD Methods.

We validate CTMD through retrospective enrichment calculations on 4 different therapeutically relevant targets, including soluble proteins (kinases) and membrane proteins (G-protein coupled receptors). The core goal of this study is to show how effectively CTMD can rank ligands after an initial virtual screen. By using datasets where pre-docked poses and *in vitro* validated hits were already available, we were able to isolate the performance of our ranking method from the common errors found in ligand pose prediction. Our results demonstrate that CTMD comprising around 3 independent metadynamics simulations of 5ns each provides statistically significant and robust enrichment over both docking and AI-based cofolding methods in differentiating chemically distinct binders from non-binders. In addition, CTMD does not suffer from any memorization habits that state-of-the-art co-folding methods are plagued with^7,16^ – the enrichment is not for physically wrong reasons. CTMD satisfies the criteria for an effective screening method: it is accurate, simple to implement - requiring minimal parameter optimization - and computationally inexpensive to perform, offering an intuitive interface for general practitioners. Through empirical studies we also find it to be remarkably robust with respect to choice of precise metadynamics parameters. Finally, it is easy to implement in any existing metadynamics based molecular dynamics code given that all we seek is calculation of the scalar quantity in Eq. 1 without any fitting or optimization. CTMD should serve as an early-stage enrichment tool for the wider pharma-ceutical and academic communities, positioned between fast but approximate docking or AI-based co-folding approaches and slower, more accurate free-energy calculations. By enabling better triage up front, it is expected to significantly reduce both cost and personnel effort across drug-discovery campaigns.

The paper is organized as follows: First, we evaluate enrichment performance on soluble proteins, using Janus Kinase 2: JAK2 (a kinase) and AmpC (*β*-lactamase) as representative systems where experimental binder/non-binder classifications and structural data are readily available. Second, we compare the performance of CTMD against traditional docking and AI/co-folding algorithms applied to G-protein coupled receptors (GPCRs), specifically the CB1R and A2*AR* receptors.

## Results and Discussion

### Soluble Protein Case Study: JAK2 & AmpC

We selected JAK2 and AmpC as representative case studies of applying CTMD for soluble proteins. AmpC is a *β*-lacatamase enzyme, which can aid bacterial antibiotic resisitence.^17^ JAK2, a kinase involved in immune mediated inflammatory diseases, represents a significant therapeutic target with well characterized small molecule inhibitors.^18^ Using the dataset published with detailed ligand pose prediction and Absolute Binding Free Energy Perturbation (ABFEP) calculations,^19^ we evaluated the accuracy of our CTMD protocol in discriminating between binders and non-binders. This JAK2 dataset contains 41 molecules, consisting of 14 binders and 27 non-binders, resulting a 34% hit rate-significantly higher than what is observed in typical in virtual screening campaigns. To create a dataset reflecting a more realistic hit rate, we performed metadynamics on all 41 molecules but randomly sub-sampled 5 binders at a time with replacement while retaining the 27 non-binders for the enrichment analysis. This 5:27 binder/non-binder ratio (≈16% hit rate) better reflects real world screening conditions where hit rates typically range from 0.1% to 5 %. Depending on the availability of ligands, we maintain a base hitrate at or below 16%. Whenever fewer ligands than available are used for the analysis, we report all results after multiple independent repeats of subsampling, each time picking a different subset of the binders to ensure statistically reproducible and bias-free metrics. We used the complete dataset for JAK2 owing to published results with ABFEP with which to compare. For all other systems, we filtered out similar ligands during our selection process (see the Section **Methods** for details.)

We evaluated performance using two complementary metrics: 1) Boltzman-Enhanced Discrimination of Receiver Operating Characteristics (BEDROC) scores,^20^ which emphasize early enrichment by heavily weighting top-ranked molecules, and 2) enrichment factors at specfic selection thresholds. Enrichment factor was defined as follows:

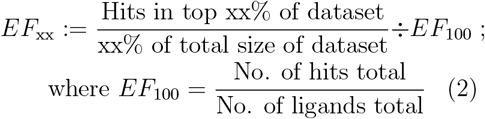

Here xx is the selection threshold indicating the top xx% of the dataset. Values *>* 1 indicate improvement over random chance.

Despite being the standard initial filter in virtual screen pipelines, docking scores are poor ranking strategies for subselecting ligands from within the top few. Indeed, for the case of JAK2, we find that GLIDE XP scores fail to pick any binders in the top 10. The relative performance of other well-optimized docking scores are highlighted in Figs. 1 (D) and 2 (B) and (D). This poor performance at the ranking stage, despite docking’s utility for the initial pose prediction, further underscores the limitations of emperical docking scores as a scoring metric after preliminary enrichment by virtual screening. Conversely, Absolute FEP calculations as reported and performed in Ref.^8^—while computationally expensive—were promising, yielding an enrichment factor of 3.5 at 20% (2.5 at 30%). However, as previously noted, the high computational cost of Absolute FEP limits its application in early enrichment studies. In contrast, CTMD demonstrates an enrichment factor of 2.5 at 30% (2.1 at 20%), which inspite of being poorer than ABFEP, has the advantage of tractably improving the success rate of future calculations. Interestingly, The AI-based co-folding method: Boltz-2, showed the strongest performance for JAK2 (BEDROC20 = 0.92, enrichment factor ≈ 4 at 20%, 3.2 at 30%). Boltz-2 achieved impressive enrichment, performing on par with or even exceeding ABFEP. However, we noted a significant overlap in chemical space, with around 57% of binders sharing a Maximum Common Substructure (MCS) of *>*90% with the training set. This high structural similarity suggests that the observed performance may be influenced by data redundancy rather than understanding the physics of the proteinligand binding.

**Figure 1:**
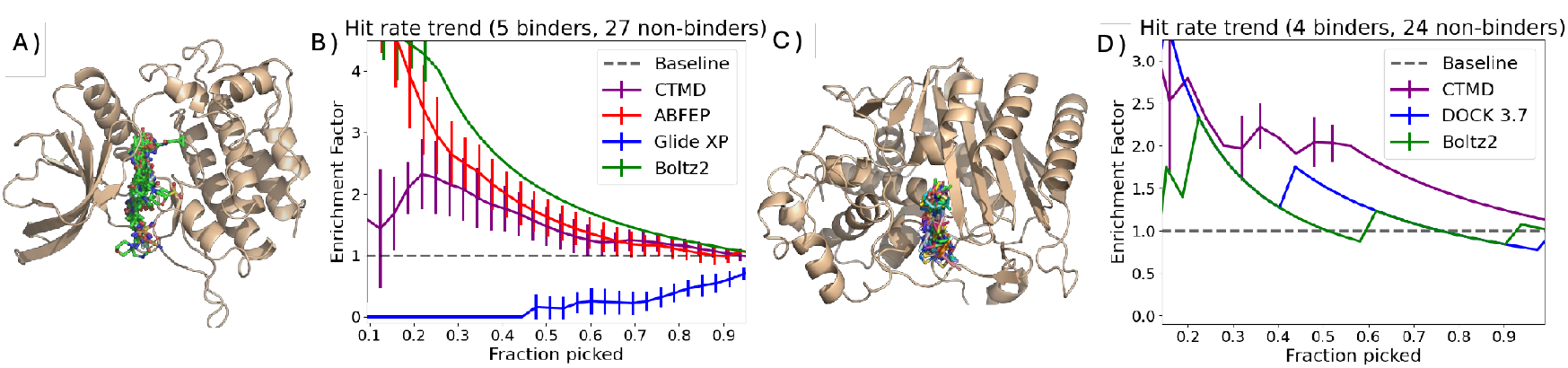
Improvement of hit rate (i.e. enrichment factor) over the docking baseline using different scoring methods for the soluble protein case studies A) JAK2 C) AmpC. B) and D) show enrichment factor (relative increase in fraction of hits from random). Docking scores (blue) usually enrich by segmenting a large database into potential binder and non-binders but are terrible at ranking true binding probability within the top few. ABFEP calculations (red) are computationally expensive but much more accurate. CTMD (purple) achieves a nice trade-off of providing some enrichment over docking with minimal parameter tuning and compute. Boltz-2 performance (green line) varies the most between both systems, a point we revisit in Fig. 3.

**Figure 2:**
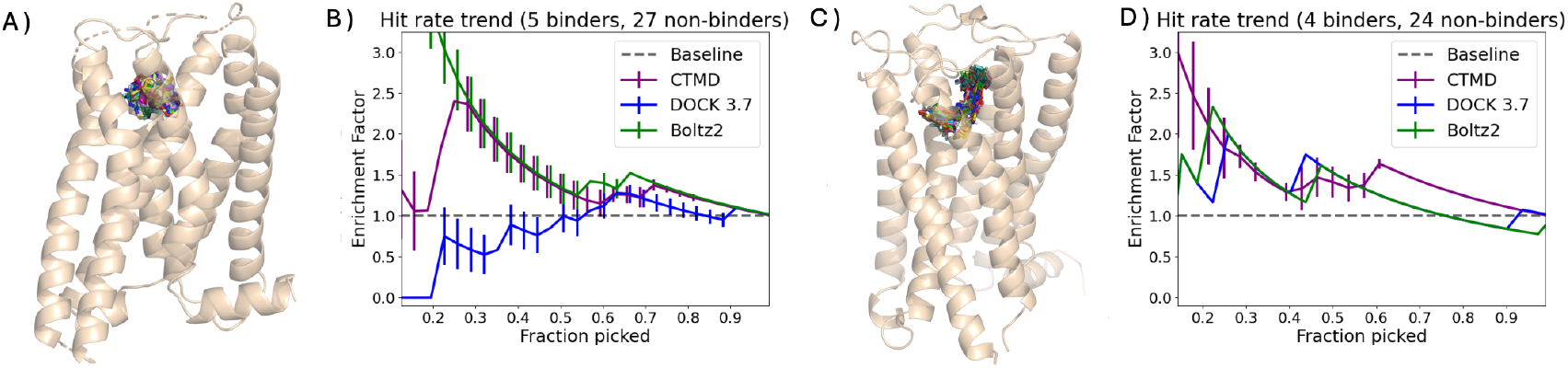
Improvement of hit rate (i.e. enrichment factor) over the docking baseline using different scoring methods for the membrane protein case studies A) Alpha2AR C) CB1R. Docking scores (blue) usually enrich by segmenting a large database into potential binder and non-binders but are terrible at ranking true binding probability within the top few. CTMD (purple) achieves a consistent performance, providing some enrichment over docking with minimal parameter tuning and compute.

In order to test CTMD’s generalizability, we applied the same protocol to AmpC, a structurally distinct *β*-lactamase enzyme. CTMD demonstrated consistent performance (enrichment factor ≈ 2.5 at 30%, Figure 1E-F), significantly outperforming docking and, notably, substantially exceeding Boltz-2, which fell below even DOCK 3.7 scores. This contrasting performance of Boltz-2 across JAK2 (excellent) and AmpC (poor) exemplifies the consistency problem facing current AI methods, whereas CTMD maintained robust enrichment across both targets.

### Membrane Protein Case Study: Alpha2AR & CB1R

To demonstrate the generalizability and robustness of the CTMD protocol across diverse protein classes, we extended our evaluation to membrane proteins—specifically G-protein coupled receptors (GPCRs). We selected the Alpha2AR adrenergic receptor (Alpha2AR) and the Cannabinoid-1 (CB1) receptor as representative membrane protein targets. Alpha2AR represents a clinically relevant non-opioid therapeutic target for pain control,^21^ while the CB1 receptor is a implicated in therapeutics ranging all the way from anxiety and pain to nausea and obesity.^22^ These systems present distinct challenges compared to soluble proteins, including the presence of lipid bilayers, different local environments, and unique conformational dynamics. Critically, we emphasize that no system-specific modifications were made to the CTMD protocol between soluble and membrane proteins beyond the inclusion of an explicit lipid bilayer and adjustment of the simulation box geometry—underscoring the method’s transferability and ease of implementation.

#### Alpha2AR Dataset and Enrichment Performance

For Alpha2AR, we utilized a dataset derived from experimental binding affinity measurements,^21^ where binders and non-binders were classified based on *pK*_*i*_ values. To establish a meaningful separation between functional classes and avoid ambiguity near the classification boundary, we applied stringent criteria: only compounds with *pK*_*i*_ *<* 5 were considered non-binders, while those with *pK*_*i*_ *>* 6 were classified as binders. This conservative approach ensured a clear pharmacological distinction between the two groups. Following our subsampling strategy to reflect realistic screening conditions, we maintained a base hit rate at or below 16%, consistent with our soluble protein benchmarks.

As shown in Figure 2A-B, CTMD demonstrated consistent enrichment performance for Alpha2AR, achieving an enrichment factor of approximately 2.5 at 30 % selection threshold. This performance significantly outpaced traditional docking methods, with DOCK 3.7 scores showing minimal enrichment beyond random selection (enrichment factor ≈ 1.0-1.2 across most selection thresholds). The CTMD protocol’s ability to discriminate binders from non-binders in this membrane protein context—without any parameter retuning—validates its robustness and highlights its practical utility for GPCR-targeted drug discovery campaigns. Notably, Boltz-2 exhibited strong performance on Alpha2AR (BEDROC=0.85 at *α* = 30, enrichment factor ≈ 2.0-2.5 at 30%), approaching or matching the enrichment provided by CTMD in this particular system. Although Boltz-2 appears to converge with CTMD’s performance in this specific system, subsequent analysis presented in the next section indicates that this high enrichment is likely a consequence of systematic memorization artifacts.

#### CB1 receptor Dataset and Enrichment Performance

The CB1 receptor dataset was unique due to significantly lower count of only 4 validated binders as compared to other systems presented in this work. While a similarity cutoff of 0.4 was sufficient for all other targets, CB1R required a more permissive threshold of 0.5 to pick these 4 binders. To establish a standardized evaluation framework, 24 experimentally validated non-binders were incorporated, resulting in a baseline hit rate of 14.2%—consistent with the rest of the study.

Fig. 2 reveals that CTMD maintained robust performance even under these challenging conditions, achieving an enrichment factor of around 1.7 at 30% selection. While this represents slightly more modest enrichment compared to the other datasets, it nonetheless demonstrates statistically significant improvement over random selection through the consistency of CTMD’s performance across multiple systems which underscores its scalability and reliability.

Boltz-2 showed reasonable enrichment for both membrane systems, significantly outperforming both DOCK scores and CTMD. However, this enrichment is brought into question when considering that Boltz-2 cannot distinguish between different membrane proteins, providing high levels of enrichment even against targets with artificially modified active sites. The impact of memorization in Boltz-2 is well summarized in Fig 3 and Fig. 6 of the SI. This suggests that its predictions are highly dependent on training set composition and may not reliably generalize to novel targets or ligand chemotypes.

**Figure 3:**
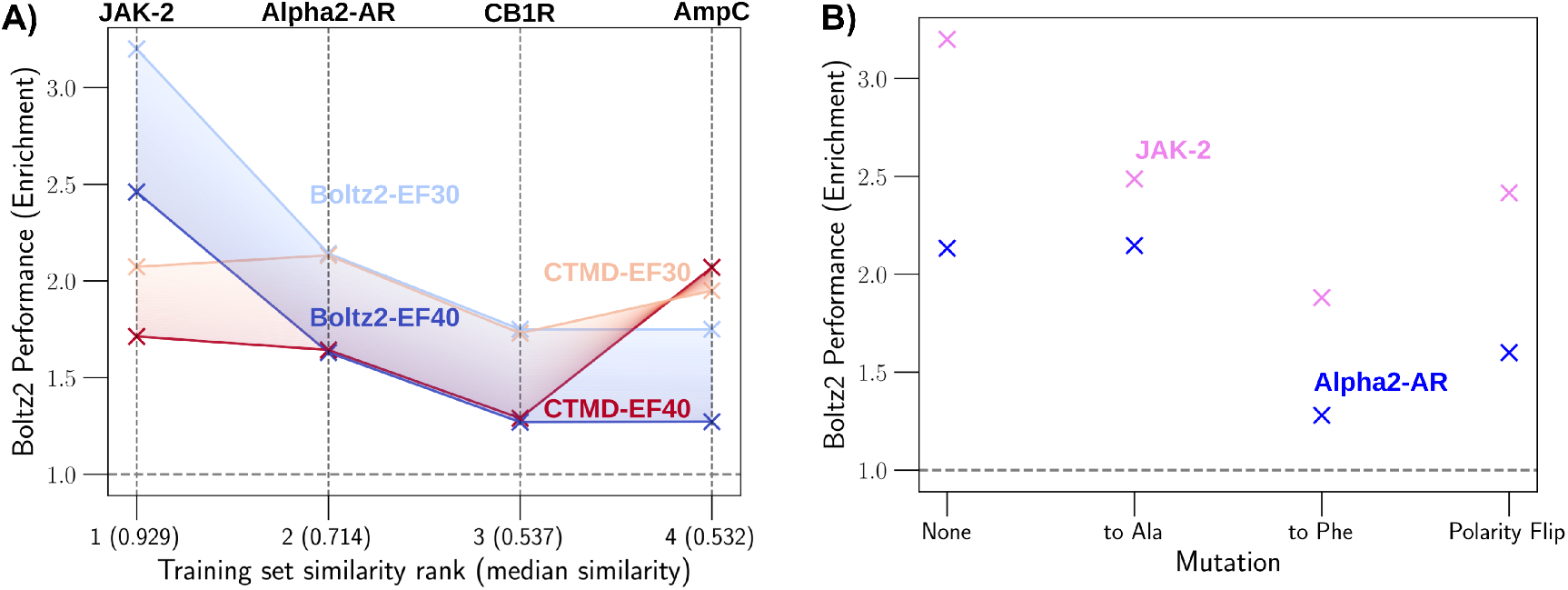
A) Boltz-2’s performance (dark and light blue curves) - as measured by enrichment factor at 30% and 40% (light and dark curve respectively) - qualitatively correlates with similarity. Systems are sorted according to median substructure similarity to the training data - value shown in parenthesis on the X-axis. CTMD (Red and pink) provides a stable enrichment across all systems (here limited to CheMBL). A summary of different mutations at protein active site (defined as the residues within a radius of 3 Å of the Co-Xtal ligand) to test Boltz-2 memorization. For each system, key active site residues were mutated in different ways i) All mutated to ALA ii) All mutated to PHE iii) Polarity and charge flipped. In all cases, the enrichment factor at 30% is consistently significantly higher than random (should be =1) for JAK2 and Alpha2AR, indicating Boltz-2’s capacity for enrichment even in the absence of a functional active site.

**Figure 4:**
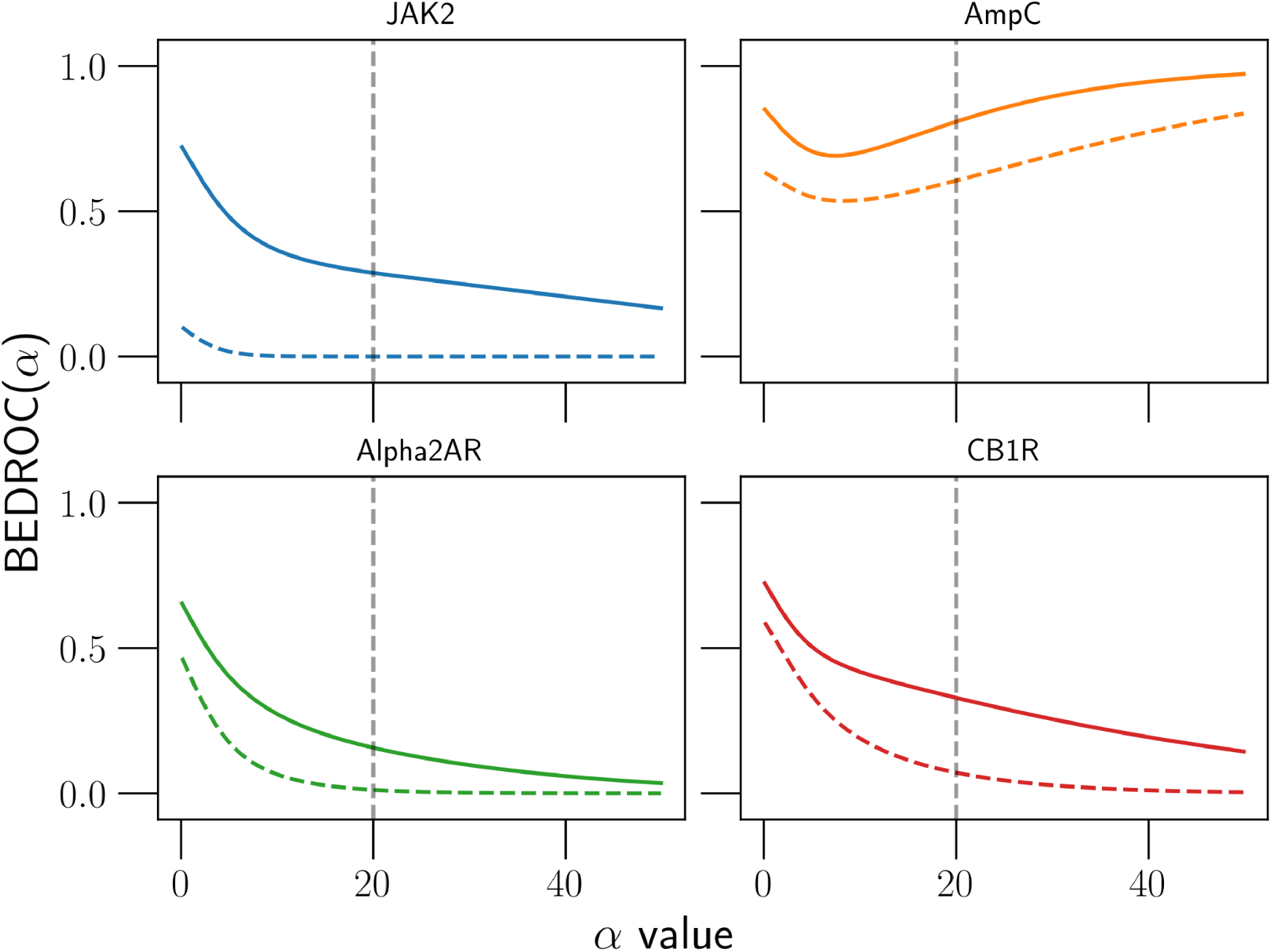
Plot of BEDROC profile as a function of *α* for each of the 4 systems tried. The solid line represents CTMD. The dashed line represents the docking score.

**Figure 5:**
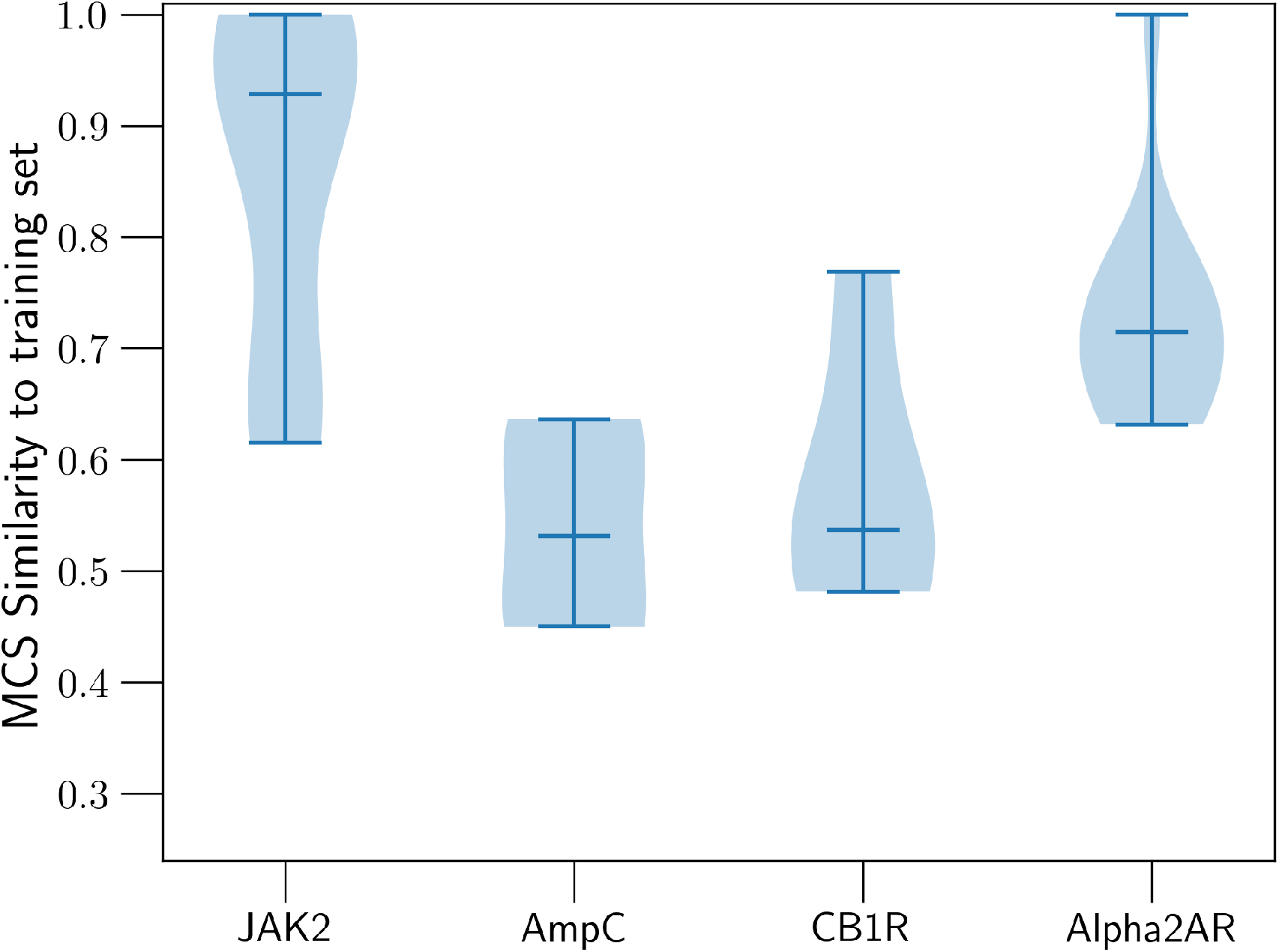
Distribution of Maxmimum common substructure (MCS) similarity of binders in our benchmark set to Boltz-2’s training data. The horizontal line inside each distribution represents the median value.

**Figure 6:**
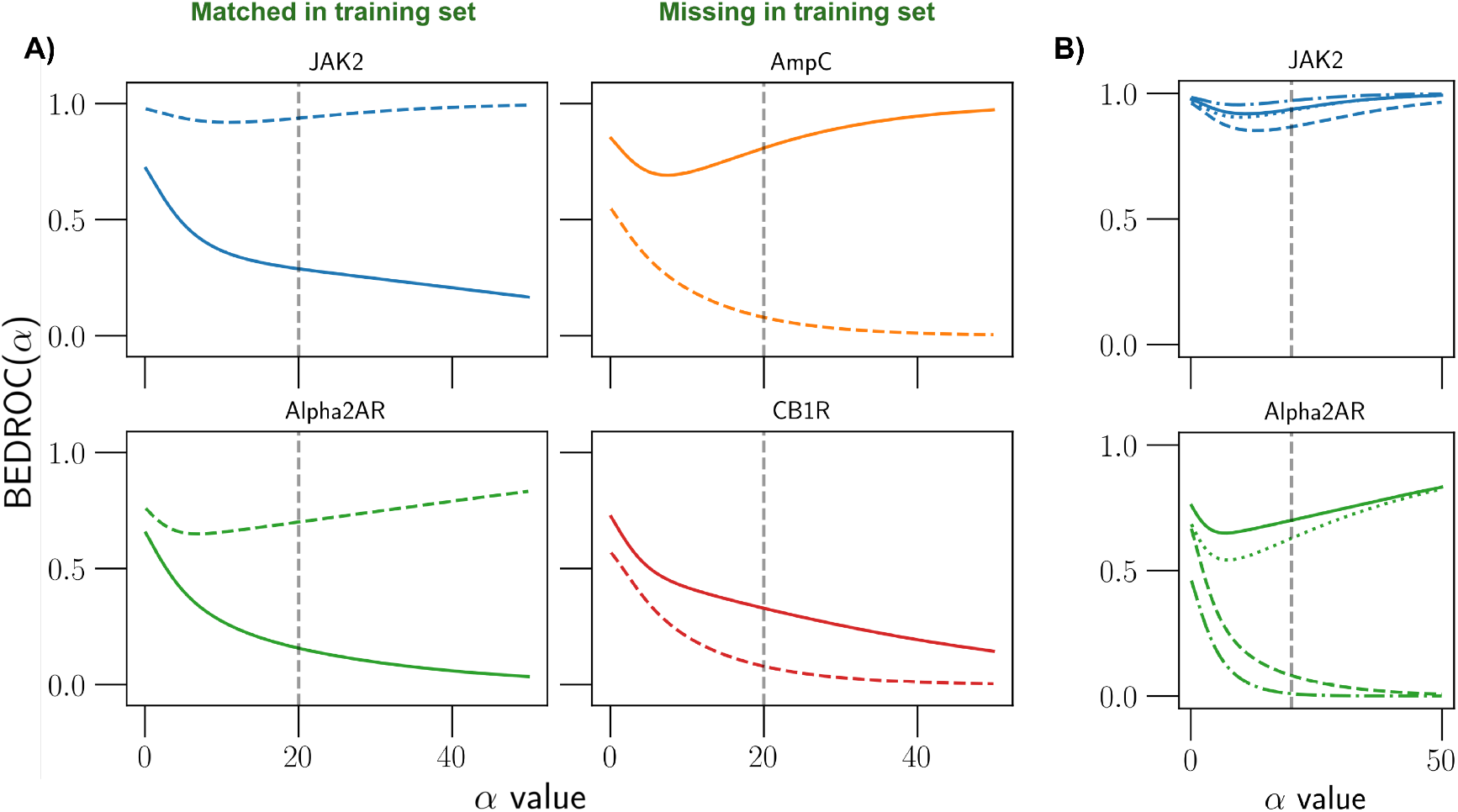
A) Plot of BEDROC profile as a function of *α* for each of the 4 systems tried. The solid line represents CTMD. The dashed line represents the Boltz-2. Notice that significant performance improvements are seen when ligands have high similarity to training set. B) Evidence of memorization in Boltz-2. Notice the early enrichment even when the active site is modified i) dashed - All to PHE; ii) dotted - All to ALA; iii) dot-dash - Flipped Polarity. The solid line is for the wild type

### Ligand-biased memorization in Boltz-2

While Boltz-2 demonstrated impressive enrichment performance on certain systems—particularly JAK2 (Fig. 1: BEDROC20 = 0.92, enrichment factor ≈ 4 at 20%) and Alpha2AR (Fig.2: enrichment factor ≈ 2.0-2.5 at 30%)—the striking inconsistency of its predictions across our test suite raised serious concerns about the reliability of AI-based co-folding methods for prospective drug discovery applications. The dramatic performance variability—from rivaling or exceeding ABFEP on JAK2 to falling below baseline docking on AmpC—suggested that Boltz-2’s apparent success on certain targets might stem from training set memorization rather than genuine physical understanding of protein-ligand interactions.

In line with previous studies^23^ that have shown that co-folding methods are prone to memorization for ligand pose prediction, here we extend such analysis to probe the generalizing abiltity of Boltz-2 to discriminate the binder vs non-binder in the context of prospective virtual screen. To investigate this hypothesis systematically, we first examined the relationship between Boltz-2’s enrichment performance and the structural similarity of test set binders to compounds in its training data. We computed the Maximum Common Substructure (MCS) similarity between our test set binders and ligands present in ChEMBL (a major component of Boltz-2’s training corpus). As shown in Fig. 3A, we observed a strong positive correlation between median MCS similarity and Boltz-2’s enrichment factor at 30% selection threshold.

However, training set overlap alone does not definitively prove memorization, as high-performing ligands for a given target may naturally share common pharmacophoric features. To rigorously distinguish between genuine physics-based scoring and memorization artifacts, we designed a series of negative control experiments. To decouple memorization from physical understanding, we subjected the JAK2 & Alpha2AR binding site to three ‘negative control’ mutations designed to abolish or invert native interactions:

- Alanine Scanning (All-ALA): All key binding site residues were mutated to alanine, effectively removing side chains and eliminating specific interactions (hydrogen bonds, *π*-stacking, electrostatics) required for ligand recognition. This mutation creates a featureless hydrophobic cavity incapable of selective binding.
- Steric Hindrance (All-PHE): All active site residues were replaced with phenylalanine, introducing bulky aromatic side chains that physically occlude the binding pocket. This mutation prevents ligand accommodation through severe steric clashes.
- Polarity/Charge Inversion (Polarity Flip): All key polar and charged residues were systematically inverted (e.g., acidic → basic, polar → hydrophobic), disrupting the electrostatic complementarity essential for binding. This creates a “mirror image” active site with reversed interaction preferences.

We reason that in each case, since we generated protein structures with ghost active sites—binding pockets that retain overall topology but lack functional interaction capability, a model that has learnt the physics of protein ligand interaction should predict that ligands cannot bind to these mutant proteins, resulting in random ranking (enrichment factor ≈ 1.0). Conversely, persistent enrichment would indicate that the model is scoring based on ligand features alone, independent of the actual protein structure—a clear signature of memorization.

The results of these negative control experiments, summarized in Fig. 3 (B), reveal profound and systematic memorization artifacts in Boltz-2. For both JAK2 and Alpha2AR, Boltz-2 demonstrated statistically significant enrichment across all three mutation types, with enrichment factors consistently exceeding random chance. In the JAK2 case, enrichment factors ranged from 1.5 to 2.5 at 30% selection threshold across all three mutation classes, with the All-ALA mutant showing enrichment factor ≈ 2.2—nearly identical to wild-type performance. Similarily for the Alpha2AR case, enrichment factors remained elevated (1.3-2.0) even for the most disruptive mutations, indicating that Boltz-2 continued to preferentially rank known binders higher than non-binders despite the absence of a functional binding site.

These findings are particularly striking for the All-PHE and polarity inversion mutations (Flip mutations), where steric occlusion or reversed electrostatics should completely abrogate binding. The persistence of enrichment under these conditions unambiguously demonstrates that Boltz-2 is scoring ligands based on intrinsic molecular properties correlated with binding in the training set, rather than evaluating protein-ligand complementarity for the specific target structure at hand. In contrast the often cited limitations of static docking protocols, function here as a crucial safeguard. Any docking protocol that enforces steric and electrostatic constraints would fail to produce any valid pose within such abrogated sites, correctly indicating that binding cannot occur (see Fig. 7 for example). Because CTMD depends on this initial docked pose, it inherits this essential physical constraint, ensuring that enrichment arises from legitimate protein ligand interactions rather than the AI-driven artifacts observed in Boltz-2. Thus by construction we expect that the equivalent of Fig. 3 (B) for CTMD is no enrichment.

**Figure 7:**
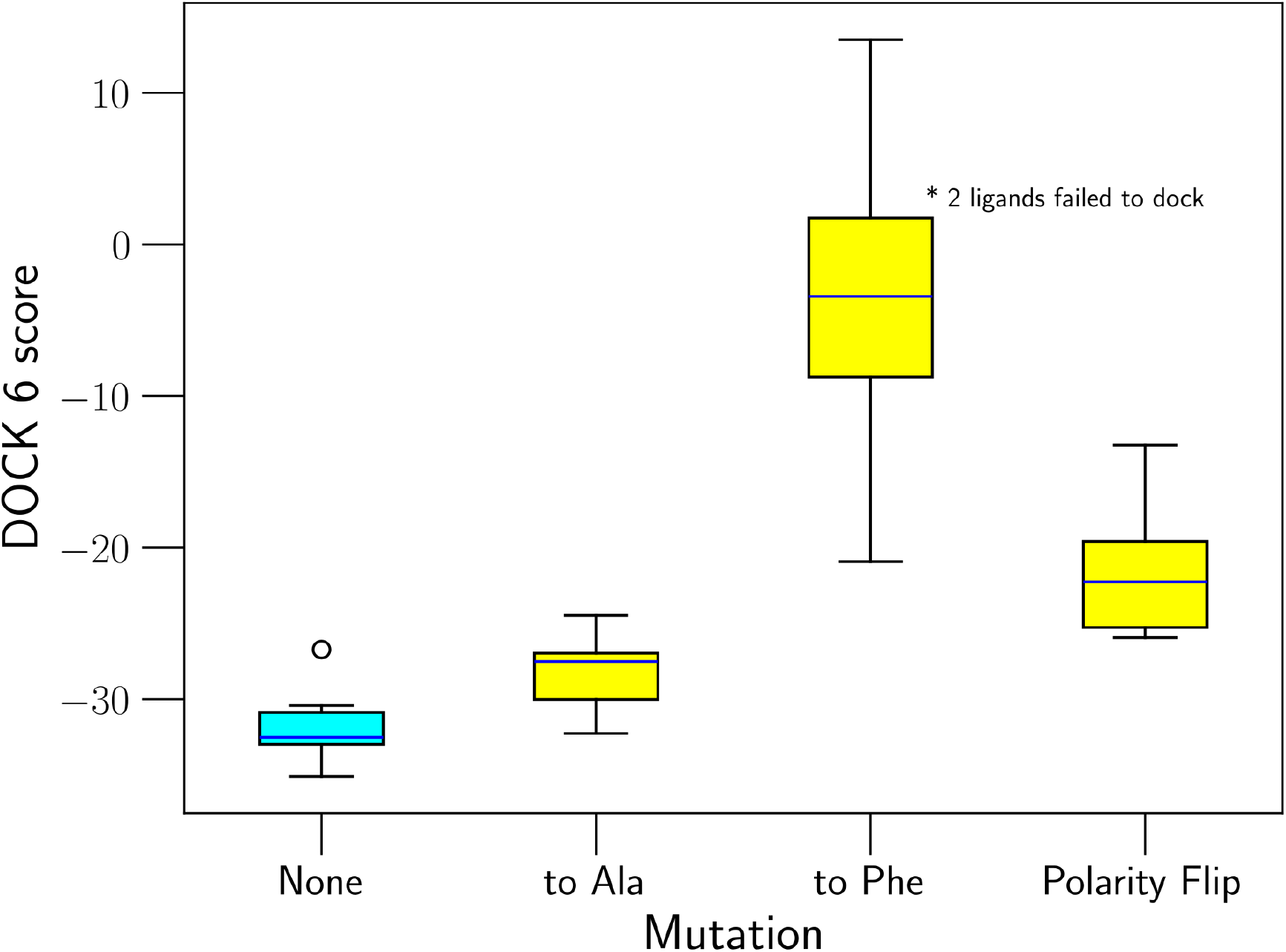
Increase in DOCK 6 score distributions when mutations to the active site are introduced. The blue horizontal lines in each box represent the medians of the distribution. The wild-type (no mutation) is shown in cyan. The mutants are shown in yellow.

## Conclusions

In summary, we introduce CTMD (c(t)-based metadynamics) as a physics-informed, highthroughput hit-triaging strategy. By using nonequilibrium reversible-work estimate *c*(*t*) as in Ref.^14^ from a small number of short, independent well-tempered metadynamics runs, CTMD avoids the overhead of fully converged free-energy calculations while still providing a robust discriminator of binding stability across chemically diverse ligands. Across a wide variety of protein systems - both soluble and membrane bound - CTMD shows an enrichment factor at 30% (EF30) of 1.8-2.0, indicating that over half the true binders will be enriched to less than a third of the full data, even over the baseline docking enrichment. CTMD is computationally inexpensive and rapid, allowing it to behave as a filter before the expensive, accurate experiments take place, greatly cutting down downstream costs. Furthermore, because CTMD enrichments do not correlate with ligand similarity to existing databases, the protocol is suited for prospective virtual screens on novel targets. This allows for the discovery of “first-in-class” binders, providing a critical advantage over AI-based methods that are often limited by their reliance on known chemical signatures. Crucially, CTMD is fast, transferable across targets with minimal tuning, and not susceptible to memorization-driven artifacts, making it realistic and practical for general users and prospective screening pipelines for unseen targets. At the same time, our results reinforce that AI-based co-folding remains far from a reliable enrichment engine for virtual screening: despite impressive structure prediction capabilities, it does not yet deliver consistent, statistically meaningful separation of binders from non-binders, and its performance directly correlates with memorization effects, as similar non-trivial enrichment factors are obtained for heavily mutated binding sites in all 4 classes of systems studied here. Here we compared CTMD with Boltz-2, but expect similar trends to hold for other co-folding softwares. While it is exciting to see development of open-source, AI-based methods for drug discovery, democratization and AI enabled advances should not happen at the cost of quality standards, including memorization of known data. The alternative is to use docking scores directly, which we have shown are too crude to successfully rank ligands top hits among themselves, which is needed for hit-triaging. Taken together, CTMD offers an immediately usable, “embarrassingly” open-source physics-based alternative for early-stage enrichment that can be implemented immediately in different drug discovery campaigns. We also believe and hope that the simple protocol we have presented here can be further optimized by us and the broad community, thereby reaching higher enrichments for the physically right reasons and not just training set rote memorization.

## Acknowledgments

This work was supported by NIH/NIGMS under Award No. R35GM142719. We thank UMD HPC’s Zaratan and NSF ACCESS (Project CHE180027P) for computational resources. P.T. is an investigator at the University of Maryland-Institute for Health Computing, which is supported by funding from Montgomery County, Maryland, and The University of Maryland Strategic Partnership: MPowering the State, a formal collaboration between the University of Maryland, College Park, and the University of Maryland, Baltimore.

## Data Availability Statement

The data and software implemented in this work will be uploaded before submission to a journal at github.com/tiwarylab

## Supplementary Information

## Methods

### Ligand selection

Each of the test systems has a very differing number of hits and decoys available to be tested. In order to standardize our experiment, we targeted a standardized hit rate of 10% where was a reasonable level to expecting from preliminary docking. We sampled as many true binders and decoys as possible ensuring a tanimoto similarity^24^ of less than 0.4 between all ligands. This usually limited most systems to less than 50 molecules.

### System selection and preparation

We hand-picked systems involving different classes of proteins from the Large-Scale Docking (LSD) database^25^ and Schrodinger’s public FEP data for JAK2 Kinase,^19^ since these databases could provide initial guesses for the protein-ligand interaction pose as would be expected from most docking screens. The proteins selected here are CB1 and Alpha2A receptors (G-protein coupled receptors), JAK2 (a kinase) and AmpC (an enzyme).

The initial protein structure was retrieved from these databases and was prepared in UCSF Chimera^26^ via Dock Prep, without changing the protonation state of any residue. This structure was saved in the default Protein Data Bank (PDB) format. For membrane proteins, CHARMM-GUI^27^ was used to prepare the initial system, exporting the final structure with AMBER parameters. The ligands were parameterized using AMBER’s GAFF2 forcefield^28^ and all components were then assembled using the leap program provided in AmberTools.^29^ Finally, Na^+^ and Cl^−^ ions were added, and the system was neutralized with the ionic concentration set to 150 mM.

### Molecular Dynamics

All simulations were performed on PMEMD patched with PLUMED version 2.9.2^30–32^ with a 2 fs time step. Simulations were performed utilizing the AMBER ff19SB force field^33^ for protein, TIP3P for water molecules^34^ and, General Amber ForceField (GAFF2)^28^ for all ligands. Temperature and pressure were kept at 303 K and 1 bar using the velocity rescale thermostat^35^ and Parrinello-Rahman barostat.^36^ The non-bonded interactions were calculated with a 10 Å cutoff, and long-range electrostatics were calculated using the particle-mesh Ewald (PME) method.^37^

Each system was first subjected to a short energy minimization for 15000 steps followed by heating it from 0K to 100K at a rate of 20K/ps and then from 100K to 303K at a rate of 2K/ps. The systems were then allowed to equilibirate for 1 ns at these temperatures with restraints (k=5 kcal/mol/Å^2^) applied to all backbone atoms and with the same restrains applied only to *C*_*α*_ atoms for another 1 ns.

For the first stage of filtering, all ligands were subjected to an unbiased MD for 5ns. Any ligands that left the active site i.e. had a pose with RMSD higher than the CTMD cutoff (described later) were automatically given a score of 0. For the ligands that remained, multiple metadynamics trajectories were fired from the final frame of the 5ns MD. The collective variable (CV) biased was protein-aligned ligand RMSD where the protein atoms were aligned and the RMSD of the ligand atoms was calculated to the reference. The reference was taken to be the start of the unbiased MD instead of the end to honour the poses predicted by docking and the heavily penalize pose drift during unbiased MD. The final system sizes are provided in Table **CTMD Simulation Summary**.

### CTMD Protocol

All Metadynamics simulations, unless explicitly mentioned used only this RMSD as the Collective Variable (CV) with a gaussian *σ* of 0.015 kJ/mol and a height of 1.75 kJ/mol, a bias factor of 10, and a pace of 1 ps and a run length of 5 ns.

Using RMSD as the CV allows biasing the ligand’s coordinates relative to the protein in a completely unsupervised manner. This ensures that our method translates between systems with minimal fine-tuning of variables - which is extremely effective for drug discovery where it is difficult to know apriori the parameters that will provide enrichment without access to known binders.

*c*(*t*) as computed in Eq. 1 serves as an excellent proxy for the total bias added, and represents the total work done through the external biasing.^14,15^ We expect this value to remain reasonably stable for a given system. The simulation is terminated if the ligand RMSD ever crosses 6Å and stays above for 200ps. Keeping in line with the general approach of using minimum work to quantify free energy differences,^14^ we repeat the unbinding metadynamics simulation 3 times, for a total of under 15 ns of biased MD, and take the lowest C(t) value. In this work, in order to achieve better statistics, we repeated the simulations 10 times, and in order to mimic a real-world scenario, all results reported are bootstrapped by repeatedly picking 3 random trajectories from these 10, and use the mean value from 250 such repeats. The error bars are a result of this process, and where applicable, subsampling of the true binders.

### CTMD Simulation Summary

The following table summarizes the the number of binders and initial system size (for MD) picked from their respective databases in each study. Many targets had far more ligands available. However, whenever possible, binders and non-binders selected were filtered to have Tanimoto similarity below 0.4 amongst themselves. Where more than 5 distinct binders were available, only 5 binders were picked randomly for mutliple iterations of bootstrapping, and this number is used in calculating the base hit-rate.

| System | Type | # of Ligands | # of Binders | # of Decoys | Base hit-rate | System Size |
| --- | --- | --- | --- | --- | --- | --- |
| **JAK-2** | Soluble (Kinase) | 41 ligands | 14 binders | 27 decoys | 15.6% | 41000 atoms |
| **AmpC** | Soluble(Enzyme) | 28 ligands | 4 binders | 24 decoys | 14.2% | 42500 atoms |
| **CB1R** | Membrane Protein (GPCR) | 28 ligands | 4 binders | 24 decoys | 14.2% | 75000 atoms |
| **A2AR** | Membrane Protein (GPCR) | 33 ligands | 6 binders | 27 decoys | 15.6% | 69000 atoms |

### Other measures of enrichment

BEDROC is a normalized version of the Robust Initial Enhancement (RIE)^38^ score where RIE weights *early* enrichment higher than the overall enrichment, which is captured by a metric like AUROC (Area Under ROC). It is defined as follows:

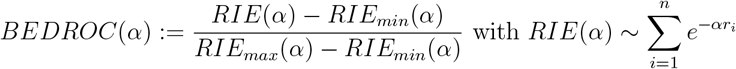

Higher values of *α* encourage earlier enrichment. It is usually plotted for different values of *α*. The literature value of *α* = 20 is commonly used to reduce the metric to one number^39^ (reported here)

#### CTMD always *improves* hit rate compared to raw docking scores

The following plot compares BEDROC profiles using just DOCK 3.7 scores for all systems (except JAK2 which uses GLIDE XP scores) compared to using CTMD ranking. In all cases, CTMD clearly improves hit-rate (independent of the value of *α* chosen as weighting factor)

#### Training set similarity of binders in our benchmarks

Many of our benchmarks have ligands which are present (including affinity scores) in the training set for Boltz-2. Needless to say, Boltz-2 performs really well on these systems. Similarity to ligands obtained from CheMBL for each of the four targets with reported affinity (IC50, EC50, Kd, or Ki) values was assumed to be reasonable information for Boltz-2 to extrapolate “hits” from. Maxmimum common substructure (MCS) fraction was used as the metric for similarity, computed using RDKit’s rdFMCS.^40^

#### Boltz-2 enrichment occasionally outperforms CTMD, but often due to memorization

The following plot compares BEDROC profiles comparing to using CTMD ranking with Boltz-2 affinity prediction. Boltz-2 is noticably unreliable, performing really well on systems with poor representation in the training data.

#### Evidence that mutations destroy the active site - DOCK 6 example

We have shown evidence of Boltz-2 memorization through examples where mutations to the protein that destroy the active site. Despite each mutant - especially ones to Phe and ones where polarity are flipped - showing a distinct drop in docking scores, Boltz-2 shows enrichment above random even with these pockets, indicating a high likelihood of memorization.

